# Robust mixture modeling reveals category-free selectivity in reward region neuronal ensembles

**DOI:** 10.1101/082636

**Authors:** Tommy C. Blanchard, Steven T. Piantadosi, Benjamin Y. Hayden

**Affiliations:** Department of Brain and Cognitive Sciences, Center for Visual Science, and Center for the Origins of Cognition University of Rochester; Department of Psychology Harvard University

## Abstract

Classification of neurons into clusters based on their response properties is an important tool for gaining insight into neural computations. However, it remains unclear to what extent neurons fall naturally into discrete functional categories. We developed a Bayesian method that models the tuning properties of neural populations as a mixture of multiple types of task-relevant response patterns. We applied this method to data from several cortical and striatal regions in economic choice tasks. In all cases, neurons fell into only two clusters: one mixed-selectivity cluster containing all task-sensitive cells and another of no selectivity (i.e. pure noise) cells. The single cluster of task-sensitive cells argues against robust categorical tuning in these areas. The no selectivity cells were unanticipated; their identification allows for improved measurement of ensemble effects. Our findings provide a valuable tool for analysis of neural data and place strong constraints on neurocomputational models of choice and control.

## INTRODUCTION

A neuron can be characterized based on its response patterns (i.e. tuning) to various task variables. These tuning properties are often used to categorize neurons into discrete groups with presumed distinct functional properties. Neurons are often selective for multiple variables at the same time; this property is known as mixed selectivity (Rigotti et al., 2013; Fusi et al., 2016; Barak et al., 2013). Mixed selectivity allows a great deal of flexibility in coding and has important implications for neural computations. It also allows neuronal ensembles to have category-free tuning (Raposo et al., 2014). Nonetheless, neurons with mixed selectivity may still fall into natural categories; for example, cells may be strongly tuned to one variable and weakly tuned to the other, or vice versa, but not in a middle state (**Figure 1**). Because measures of neural activity are noisy, it can be difficult in practice to distinguish selectivity drawn from a continuum from selectivity drawn from categorical clusters.

Understanding the distribution of tuning functions is important for several reasons. First, delineation of cell types is an important basic description of the nature of responses in a brain area. Classification provides a useful way to simplify what are often complex and heterogeneous response patterns. Second, it provides important constraints on neural models. For example, important models of economic choice rely on classification of cells into discrete types based on selectivity for particular stimuli (Soltani et al., 2006; Hunt et al., 2015; Rustichini and Padoa-Schioppa, 2015; Chau et al., 2014; Louie et al., 2011). Finally, categorization of neurons provides a possible baseline for establishing functional implications of anatomical and genetic methods that delineate specific cell types (e.g. Kvitsiani et al., 2013).

Although many types of categorizations of neural populations have been proposed in other prefrontal regions, none have been well established. Category proposals often rely on visual inspection or on methods like k-means, which produce best clusterings for an arbitrarily chosen number of clusters, but do not provide information about whether such clusters are statistically justified. Another common approach is to use the standard null-hypothesis testing cutoff (tests against p<0.05) to divide cells into categories and use the resulting cell classifications to make inferences about category function. This method can be susceptible to both Type I and Type II errors (Maxwell and Delaney, 1993). Three aspects of this method are particularly problematic. First, it throws a way a great deal of information. Second, whether a neuron reaches significance will depend on the number of trials collected; this arbitrary factor will further increase variance in our estimate of the true clusterings. Third, due to the mathematics of joint probability, when data are noise limited (almost always), fewer neurons will reach significance for two variables than for either one. This bias will lead to a systematic overestimate of coding disjunctions – that is, neurons will appear more purely categorical than they really are.

The question of whether there are natural functional categories of cells in most brain regions remains unanswered. This problem is particularly salient now. Recent successes of population analysis methods have demonstrated the power of ensemble coding (Mante et al., 2013; Stokes et al., 2013; Churchland et al., 2012; Kristan and Shaw, 1997; Ma et al., 2014; Hyman et al., 2012). Moerover, theoretical advances confirm the flexibility and power of mixed encoding schemes (Rigotti et al., 2010; Rigotti et al., 2013; Ganguli and Sompolinsky, 2012; Pouget and Sejnowsky, 1997; Fusi et al., 2016; Barak et al., 2013). Finally, a recent high-profile study specifically investigating the question of tuning categories finds not just mixed selectivity but no evidence of categorical organization in rodent parietal cortex (Raposo et al., 2014). This finding raises the possibility that category-free selectivity may be a general property of neural systems.

We developed a novel analysis approach by using a generative Bayesian model. Our model makes statistical assumptions about the components likely to have given rise to the observed firing patterns. It identifies three potential components in any data set: 1) A *no selectivity component* that generates neurons that have no tuning to any task variables, 2) a set of *pure selectivity components* that generates neurons that have tuning to only one task variable, and 3) a *mixed selectivity component* that generates neurons that have tuning to multiple task variables. The statistical analysis assumes that the observed data come from a weighted mixture of these components and infers the weights from the neural firing patterns. Thus, it might discover that most neurons are purely selective (strong categorical organization, **Figure 1A**)or come from one large category of mixed selective cells (**Figure 1B**), or even that they are non-selective (not task relevant, **Figure 1C**). It could also show a mixture of these sets, involving multiple discrete categories (**Figure 1D and E**). Alternatively—and in contrast to clustering methods—the analysis might show that the data are not highly informative about the classification, telling us that additional techniques or larger data sets are required.

To identify these components, we use Bayesian tools (specifically Markov chain Monte Carlo - MCMC - sampling) in the programming language Stan to infer likely values of the mixture weights and parameters of these components (Carpenter et al., 2015). Our analysis starts from relatively unbiased assumptions about what the parameter values may be (see Methods). Several choices in our modeling setup are designed to make this analysis robust, in particular to outliers and the presence of no-selectivity neurons. These include deliberate inclusion of a noise component to remove the influence of no selectivity neurons, use of t-distributions as mixture components due to their robustness to outliers (Peel and McLachlan, 2000), and inclusion of variance and covariance from the underlying regression to correctly handle uncertainty in each neuron’s response profile. Our method is publicly and freely available (Blanchard, 2016).

To validate this method, we applied it to synthetic datasets with known category structure and show that it accurately infers their parameters. We then examined functional properties of neurons in four reward-sensitive frontostriatal brain areas in a variety of economic decision-making tasks (orbitofrontal cortex, OFC, Blanchard et al., 2015; dorsal anterior cingulate cortex, dACC, Blanchard et al., 2014 and Azab and Hayden, 2016; ventromedial prefrontal cortex, vmPFC, Strait et al., 2014; and ventral striatum, VS, Strait et al., 2015). For all twenty sets of variables we examined (ones relevant to economic choice and executive control) we found the same pattern: neurons fell into only two categories, one category containing all task-sensitive cells, and another one containing cells best classified as pure no-selectivity. We found no evidence for distinct categorical tuning in any of the datasets we analyzed.

**Figure 1.**
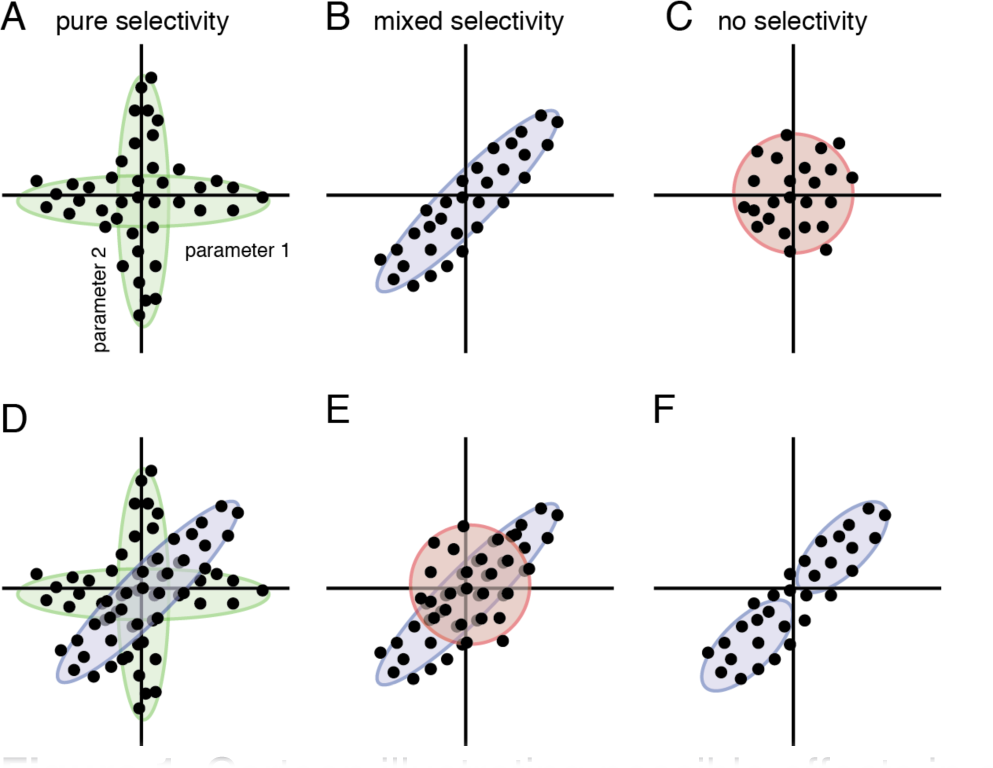
Cartoon illustrating possible effects in a dataset. Consider a set of neurons in which cells’ tuning properties for two parameters are independently assessed and then z-score transformed. Scatter plots are cartoon versions; each dot corresponds to a neuron; the two dimensions are two tuning dimensions. **A.** Neurons may fall into pure-selectivity clusters. In the example, neurons are strongly tuned for parameter 1 or 2 but not both. **B.** Another possibility is that neurons fall into a single larger cluster selective for both variables. In the presence of noise, it is often difficult to distinguish pure from mixed selectivity **C.** A third possibility is that neurons will have no selectivity for either variable, and will be best classified as non-selective. Some cells may be significantly tuned, but thus number is no greater than the false positive rate expected by chance. **D.** A population of neurons can also contain a combination of subsets that are pure and mixed, as for example when a population contains subpopulations correspond to each of two variables and a third that integrates them. **E.** Neuronal populations can also contain other mixtures of these. In particular, our data suggest that a combination of mixed and no selectivity populations are common, perhaps even universal, in reward regions. **F.** Conventional clustering methods make it difficult to judge categorical structure of populations. For example, a method like k-means clustering will divide cells into two categories (two blue ovals) even if they are statistically likely.

## RESULTS

### Description of the model

Consider an experiment in which two variables are manipulated independently across several trials. In a risky choice, for example, these variables could be reward amount and reward probability. Neural firing rates are recorded for a set of neurons over a few hundred trials as these variables vary independently. We wish to know how the ensemble encodes these two variables.

Our model employs a two-part estimation scheme. First, we z-score the firing rates; then we analyze each neuron’s responses with a linear model and compute regression weights (i.e. beta coefficients or values) for each variable as well as the variance and covariance in these estimates. The beta values resulting from these computations describe the response of each neuron to each variable while controlling for the other variable or variables, as well as possible confounding variables. Our analysis uses a separate regression from the mixture model for reasons of computational efficiency, although it is possible in principle to integrate both aspects into a single hierarchical analysis.

Thus, the basic starting point for our analysis can be viewed as a *scatter plot,* with a point for each neuron located at its x-location the estimated effect for one task variable and its y-location the estimated effect on the neuron for another (See, for example, **Figure 2**). The question we address is: what unobserved components drive the pattern in this scatter plot—are there *underlying populations* of neurons that are selective to both task variables, to one, or to none?

To answer this question, the collection of beta weights is modeled as a mixture of non-selective vs. task selective cells, and then as cell subtypes within the task-relevant categorization. The no selectivity cells are assumed to have beta weights drawn from zero. In practice, of course, the measured numerical value will be different than zero due to statistical noise in sampling and estimation; our model takes this into account. Cells that are unlikely to be no selectivity are modeled as either pure selectivity (beta is zero on one variable and nonzero on the others) or mixed selectivity (the betas come from a covariance matrix).

All of the parameters are jointly inferred from the firing patterns, meaning that we can provide estimates of the proportion of cells that are non-selective (alpha), the proportion that are purely selective (beta), the covariance matrix of cells that have mixed-selectivity (1-beta), as well as the probability that any individual cell belongs to any of these categories.

The inferred distribution on covariance matrices for the mixed-selectivity cells is particularly interesting because it tells us whether, for instance, cells that respond positively to reward probability likely also respond positively (positive correlation) or negatively (negative correlation) to increases in reward. Or, if this covariance matrix has zero covariance, that would indicate that individual cells have statistically reliable but independent responses to these two task variables.

The model also includes parameters for the “scale” of firing patterns, fit separately to each mixture component. These parameters allow the model to fit the measurement scale used empirically as well as the range of possible firing rates observed.

This feature is important because it lets the model deal gracefully with large differences in tuning strengths in different data sets or across different neuron types.

Note that the inference scheme is not dependent on a specific procedure of fitting or clustering. Our procedure is rather a statistical technique that gives the optimal (relative to our assumptions) beliefs about what parameter values are likely, given the observed data. For example, instead of simply being able to state that 40% of cells are task relevant, we can say that there is a 95% chance that between 35% and 50% of the cells are task relevant. (We provide median estimates and credible intervals below). Thus, if the data are under-informative (e.g. due to a low number of cells or trials), our technique could tell us so—the parameter values will have a large range of possible values. Moreover, due to our use of Bayesian inference techniques, the uncertainty in each parameter appropriately adjusts for the uncertainty in the others, meaning that our estimates of variables like the covariance in task selective cells correctly adjusts for uncertainty in the categorization, as well as the uncertainty in the estimated regression coefficients.

### Testing the model on simulated data

To confirm that our model works as anticipated, we first tested it on simulated data. We therefore constructed ersatz neuronal datasets. For each neuron, we generated a ‘true’ tuning parameter by sampling from a distribution that could be described as one of the three classes (mixed-tuning, pure-tuning, or no-tuning). Separately for each simulated neuron, we generated simulated trials by generating a random value for each variable for each trial, and generated neural firing as a product of the variables and the neurons’s tuning parameters (plus Gaussian noise).

We chose the strength of the tuning parameters and the amount of noise so that the regression coefficient values and variances roughly match the levels we saw in our real data sets (median absolute regression coefficient value in simulated data=0.092, range in true data sets=[0.039 to 0.132]; median regression coefficient variance in simulated data set=0.0019, range in true data sets=[0.0019 to 0.0107]). As described above, we regressed the firing rates onto their tuning variables, separately for each neuron, and then obtained estimates of regression coefficient and covariance. We then fit our Bayesian model to these outputs.

We first tested if the model could correctly identify that a population of neurons composed of a single population with mixed selectivity (**Figure 2**). The model correctly places all of its weight on the mixed selectivity component (median mixed selectivity weight=1, 95% credible interval= [0.93 to 1]).

**Figure 2.**
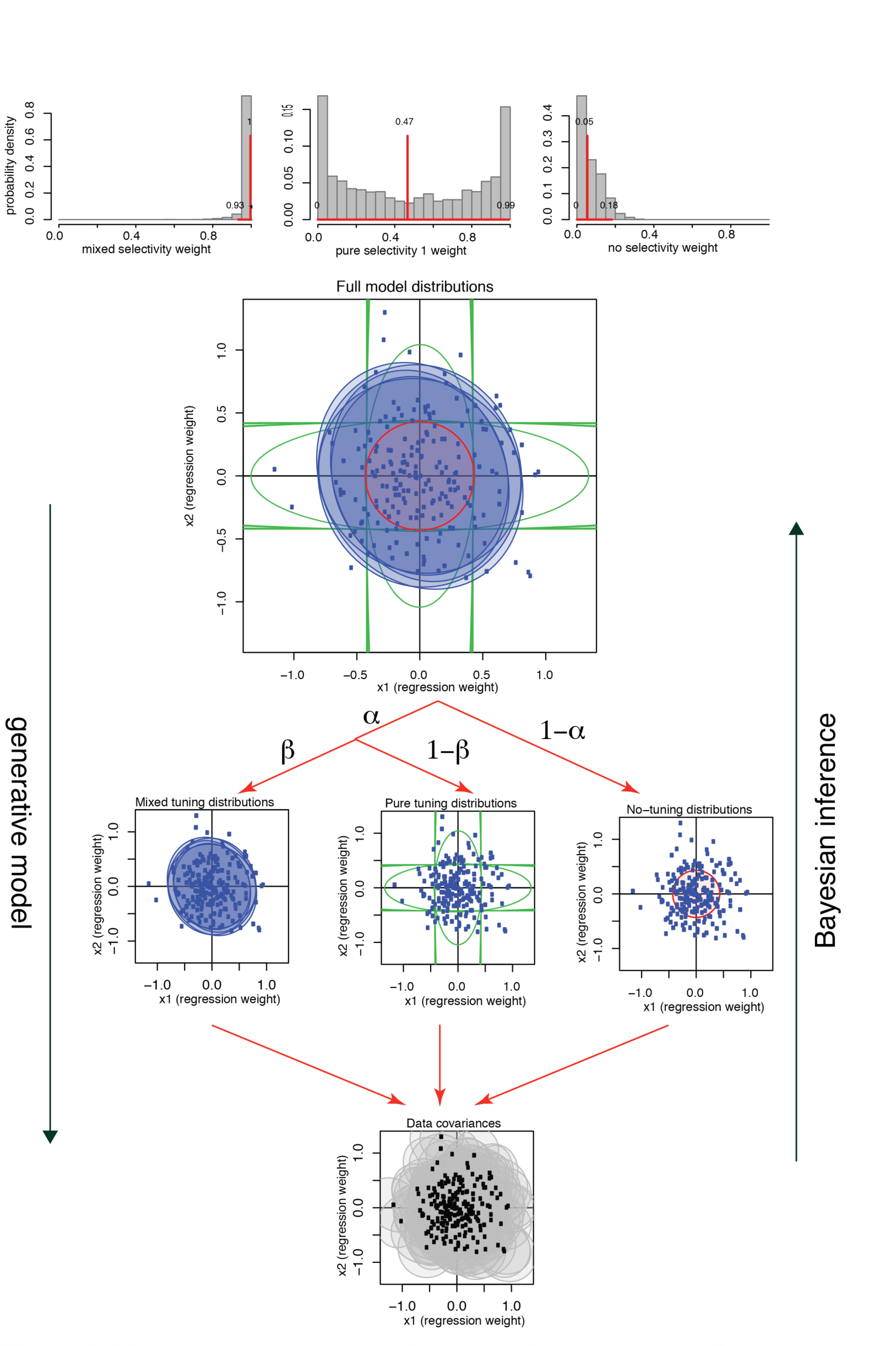
Model visualization for a mixed selectivity population. Top row shows the posterior estimates for the parameters of the model. The first scatter plot panel shows the covariance of each neuron’s tuning estimate. The following row of three scatter plot panels shows the shape of each component of the model; the five ovals come from five different iterations and their shading is proportional to the amount of weight on that component. The bottom panel shows the components combined.

We next extended our test to consider the case of a population of cells with neurons from each component (**Figure 3A-B**). In this constructed population, 25% of cells were purely selective for variable 1, 25% purely selective for variable 2, 25% had mixed selectivity for both variables, and 25% had no-selectivity to either. Our model was able to accurately estimate the proportions of no-selectivity neurons (correct answer: 0.25, model median weight: 0.25, 95% credible interval: [0.15 to 0.36]). It also captured the proportion of the signal weight that was mixed-selective (correct answer: 0.33, model median 0.4, 95% credible interval: [0.24 to 0.59]). Finally, it accurately estimated the proportion of the pure-selective neurons that were sensitive to each variable (correct answer=0.5, model median weight=0.53, 95% credible interval=[0.36 to 0.72]).

We then tested the model’s ability to detect correlations in the neurons’ tunings. In a population of neurons with mixed tuning to two variables, there is often a fixed relationship between how they are tuned to each of the two variables. For example, neurons positively tuned to one variable may more frequently be positively tuned to a second variable. This relationship would be observed, for example, if neurons encode value rather than its components (e.g. reward amount and reward probability, Strait et al., 2014; Strait et al., 2015).

To see whether our model can detect this relationship, we tested two more simulated datasets, with the same proportions of neurons coming from each of the components as above, but with a positive correlation of 0.5 between the tunings for the two variables for the mixed-selective neurons (**Figure 3C-F**). The model again was able to correctly estimate the weights for each component. It was also able to correctly estimate the correlation in the mixed-selectivity component (correct answer=0.5, median for the population of only mixed-selective neurons=0.45, 95% credible interval=[0.33 to 0.56]; median correlation estimated for population with all components=0.54, 95% credible interval = [0.33 to 0.76]).

**Figure 3.**
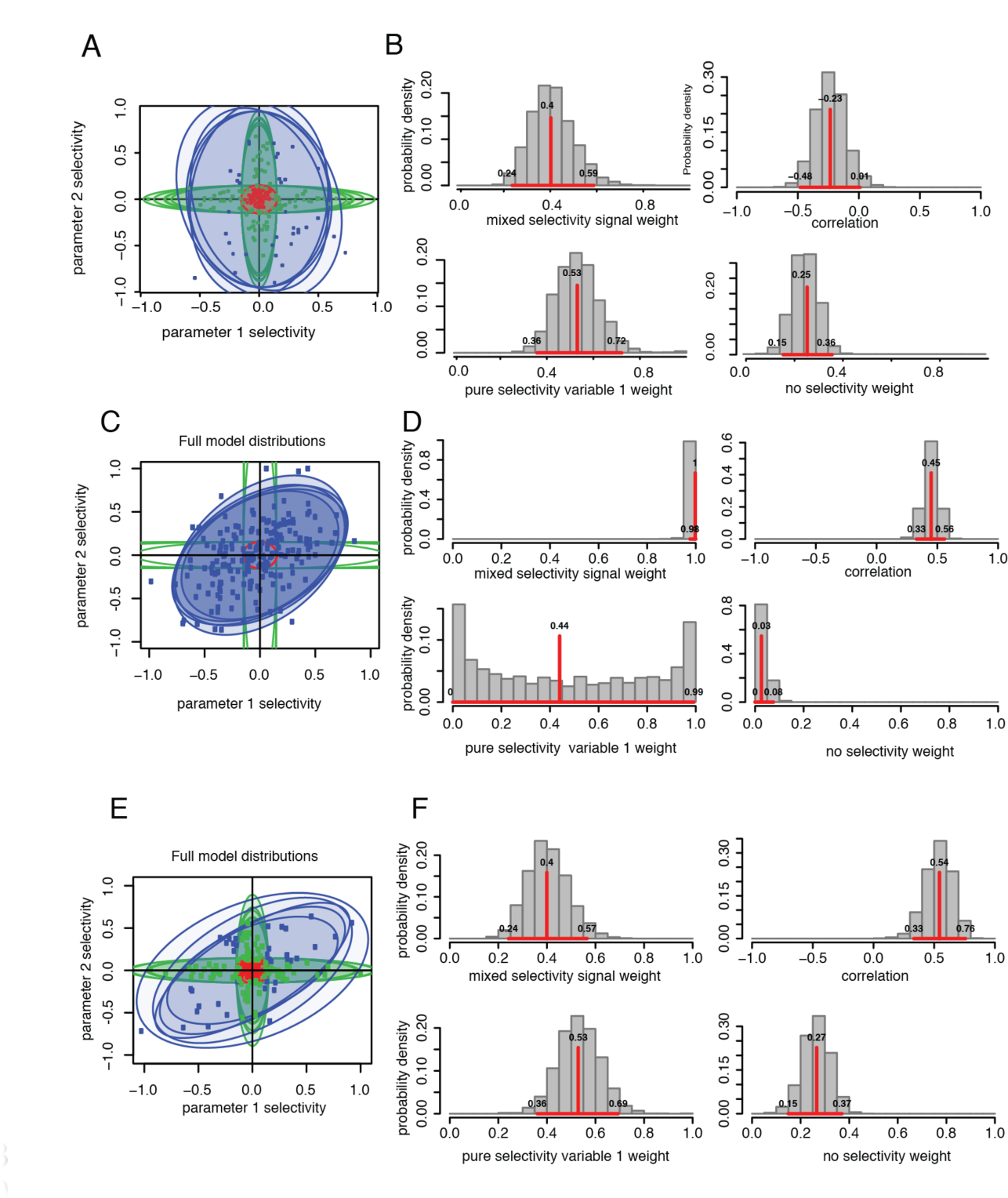
Model fit to simulated data with correlated tunings. **A.** Model visualization for simulated data with a mixed-tuned population with a 0.5 correlation between the tuning for X1 and X2. **B.** Probability distribution functions of posteriors for A. **C-D.** Same as A-B, but with a mixed and non-selective set. **E-F.** Same as A-B, but with the population split between no-tuning, pure-tuning, and mixed-tuning.

Because our model is designed to deal with extremely noisy datasets, we wanted to ensure that it converged to a single set of parameters as more data were added. We therefore tested the model’s behavior for a population of 50% mixed-selectivity and 50% no-selectivity neurons (scatter plots not shown). The weights assigned to these guesses rapidly converged as more neurons were added (**Figures 4A**). We also examined a population of 50% pure-selectivity and 50% no-selectivity neurons (**Figures 4B**). Finally, we tested a population of 25% pure-selectivity to variable 1, 25% pure-selectivity to variable 2, 25% mixed-selectivity, and 25% with no selectivity (**Figure 4C**). In all three cases, the model quickly converged to the correct weightings after observing data from between 50 and 100 neurons. Additional neurons produced greater convergence but did not qualitatively affect the data.

**Figure 4.**
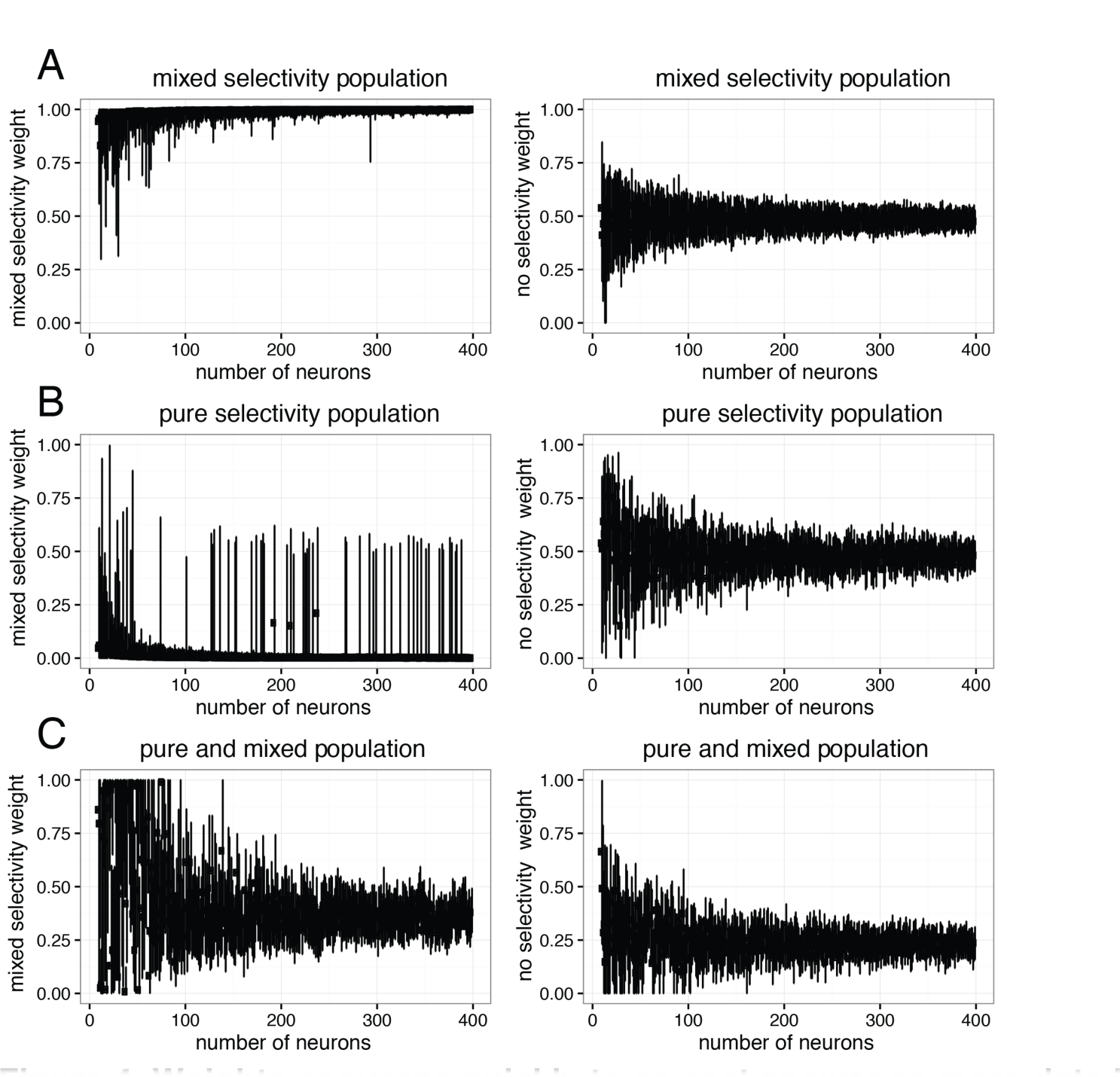
Weights converge quickly to correct answer as more data is given. Mixed selectivity (left) and no selectivity (right) weights for a mixed/no selectivity population (**A**), a pure/no selectivity population (**B**), and a mixed/pure/no selectivity population (**C**).

### Strong evidence against categorical selectivity in orbitofrontal cortex

We next applied this analysis technique to real data. We started with a dataset collected in the OFC (Blanchard et al., 2015). In the *curiosity tradeoff task* monkeys chose between offers that differ in two discrete dimensions, reward (water amount) and informativeness (information about the upcoming outcome of a gamble), which is shown by behavior to be valued, presumably because it sates curiosity (Kidd and Hayden, 2015). These two dimensions were both encoded by neurons in OFC during the offer period of the task.

Our model can answer some key questions our earlier study could do only crudely. For example, are the neurons that encode reward different from the ones that encode informativeness? Or are these variables distributed in two sets of neurons – or at random across neurons? And, are the neurons whose firing rates appeared to be unrelated to these variables simply too weakly tuned to detect an effect, or is there a way to confidently classify them as no-selectivity neurons?

The model produced clear results. We found, first, that reward-sensitive neurons and informativeness sensitive neurons do not constitute different sets of cells (**Figure 5**). Instead, they came from a single larger class of task-relevant cells with mixed tunings (Median mixed-tuning signal weight=1, 95% credible interval=[0.96 to 1]). Although the two sets are not distinct, the correlation between the coefficients is not significant (median r=0.1, credible interval=[−0.22 to 0.41]).

We did see a clear split between the task-selective cells and the no-selectivity cells (median no selectivity weight=0.43; 95% credible interval=[0.23 to 0.63]). Thus it does not appear to be the case that untuned cells are simply tuned ones for which we did not collect enough data – the model tells us that we had enough data to see an effect had it been there, at least in a substantial number of these cells. This result was not reported in that paper (nor could it have been, since the methods could not detect it).

Note that our new results, which rely on Bayesian statistics, are qualitatively different from those derived from conventional statistics because they permit inferences that take into account our uncertainty over whether or not a neuron is task-relevant. In methods that attempt to classify neurons based on a statistical threshold, the evidence provided by many sub-threshold neurons (e.g. those that do not differ significantly from zero) has no effect; in this analysis, the joint inference of each neuron’s classification and relevance uses as much information from the data as possible, making the analysis potentially much more sensitive.

Another important question in OFC is whether neurons are selective for the identity of specific offers (a labeled line code), or whether they use a single consistent format to encode the values of each of the two offers. Labeled line codes are important in many models of economic choice (Soltani et al., 2006; Chau et al., 2014; Rustichini and Padoa-Schioppa, 2015; Hunt et al., 2014; Louie et al., 2011) but recent work suggests they may not be an accurate description of neurons in decision-making structures like OFC/vmPFC (Rich and Wallis, 2016; Strait et al., 2014; Blanchard et al., 2015) and parietal cortex (Raposo et al., 2015). We first examined whether OFC neurons show categorical tuning based on side (neurons coding the left offer and neurons coding the right offer), as in LIP (Gold and Shadlen, 2007; Platt and Glimcher, 1999). Value here was defined based on the offered reward size, Blanchard et al., 2015.

Our analysis provides strong evidence against categorical selectivity (median mixed-tuning weight=1, 95% credible interval=[0.94 to 1]). Strength of tuning for left and for right offers is highly positively correlated (median correlation=0.71, 95% credible interval=[0.37 to 0.95]). Another related possibility would be that neurons are selective for the value of the first and second offer, respectively. Our results argue strongly against categorization by side as well (median mixed-tuning weight=0.98, 95% credible interval=[0.87 to 1]). Instead, neurons that are responsive to the value of the first offer are much more likely to be the ones responsive to the value of the second offer. A different possibility would be neurons sensitive to first offer value and chosen value; we see no evidence for such a distinction in OFC (median mixed-tuning weight=0.97, 95% credible interval=[0.92 to 1]). In other words, the neurons were consistently responding to the values of the offer currently presented whether it appeared first or second, the left or the right, or the offer or the chosen. The evidence points against a labeled line coding scheme.

**Figure 5.**
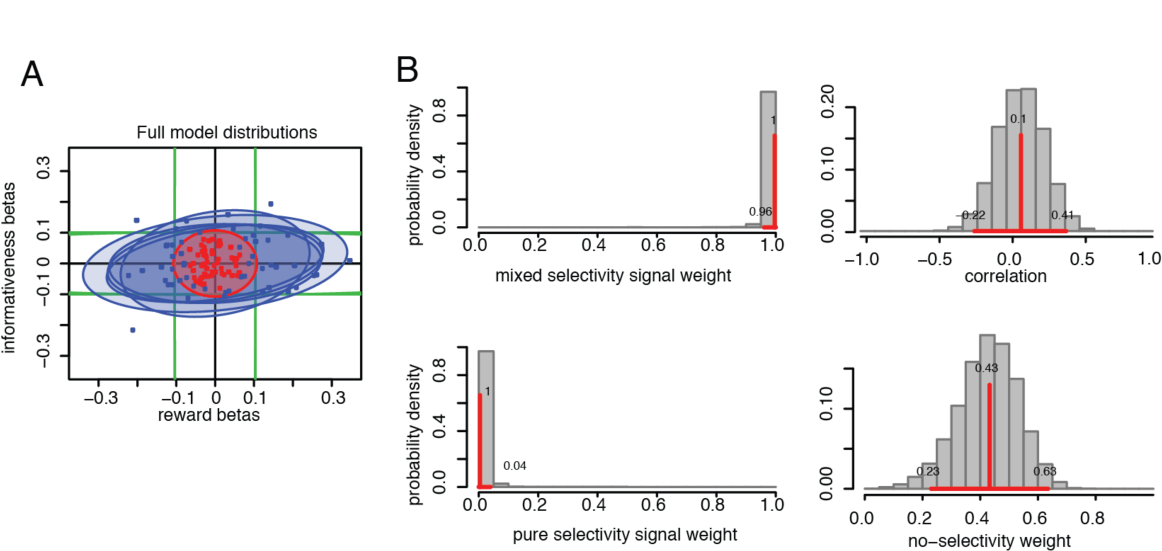
Model fit to OFC dataset 1 (curiosity tradeoff task). **A.** We fit regression weights for reward sensitivity (x-axis) and informativeness (y-axis). Cells were categorized either as mixed tuning (blue oval) or as non-selective (red oval). **B.** Posteriors. Most weight was put onto the mixed-tuning class, indicating neurons selectivity for both variables. The correlation between these variables was not found to be significant; it overlapped with zero. Pure selectivity signal weight applied to few cells. Data fit to no-tuning category shows a discrete cluster of cells that were not sensitive to either variable.

### Strong evidence against categorical selectivity in other datasets

We then considered a total of twenty pairs of variables coming from five datasets from four different brain regions (**Table 1**). We looked at firing rates of neurons in vmPFC and VS in a gambling task (Strait et al., 2014, Strait et al., 2015), dACC in a diet selection task (Blanchard and Hayden, 2014; Blanchard et al., 2015) and in a token gambling task (Strait et al., 2016; Azab and Hayden, 2016), and OFC neurons in a riskless choice task (Wang et al., unpublished data). In all cases, monkeys made economic decisions based on combining different, orthogonal, dimensions. (There were several differences in the tasks as well, see Experimental Methods for details).

In all cases, we saw the same basic pattern: neurons are categorized into two sets, a single task-sensitive set and a no-selectivity set (**Figure 6**). The task-sensitive set consisted of neurons sensitive to all task factors and the no-selectivity set was, as far as we could tell, not modulated by the variables we chose. We did not see any pairs of variables for which there are clear neuronal classes. Indeed, for all of our datasets values of over 0.05 for the pure-selective component fell outside of the 95% credible interval, meaning that with p<0.05 confidence we can reject more than 5% of neurons being purely selective (**Figure 6B**).

**Figure 6.**
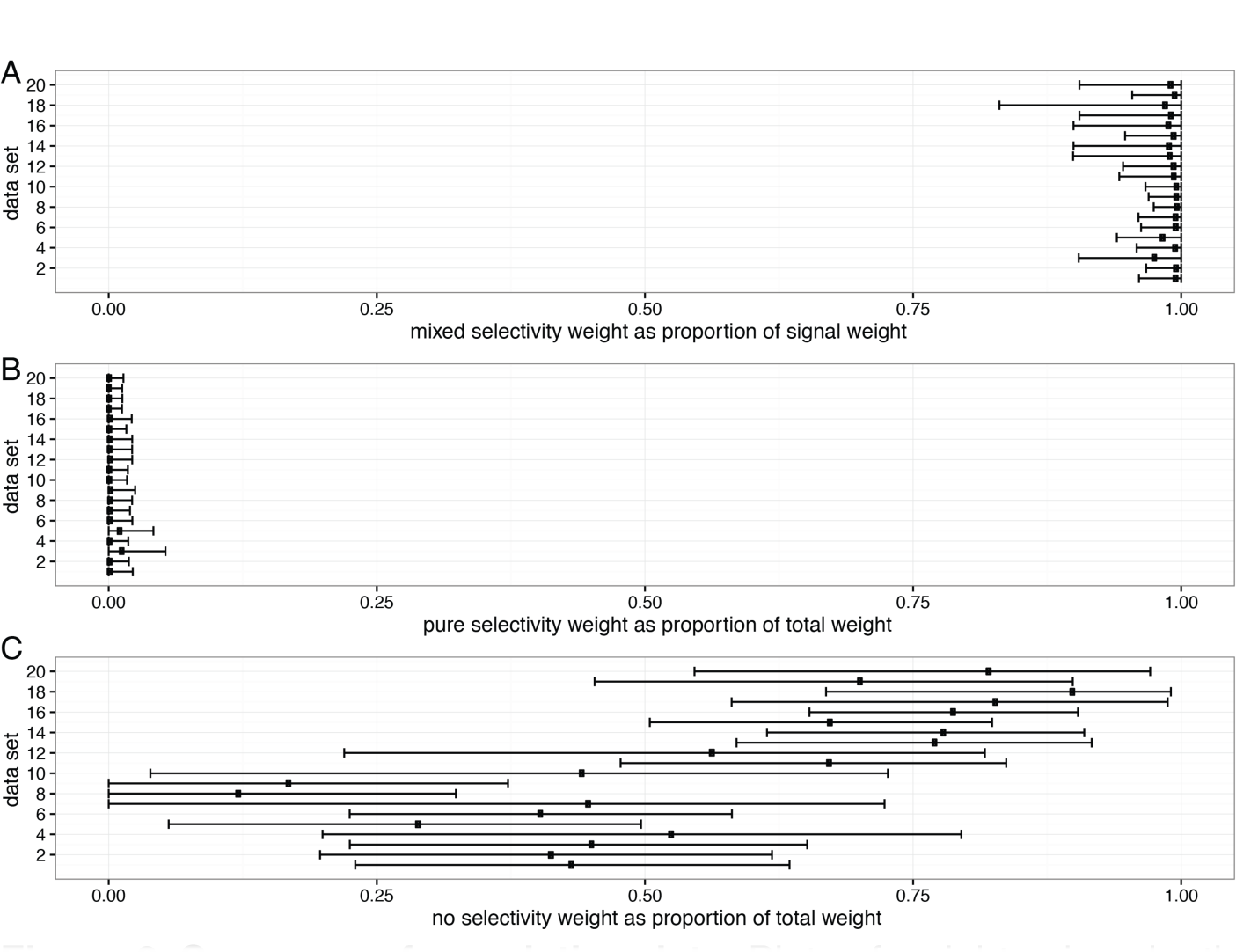
Summary of population data. Plots of weights given by the model to each of twenty neural data set (see Table 1). Bars indicate credible intervals. **A.** Weights given to mixed selectivity are high and overlap with 1.0 in all cases. **B.** Weights given to pure selectivity are weak and overlap with zero in all cases. **C.** Weights given to no selectivity are positive and do not include 0 in most cases.

**Table 1.**
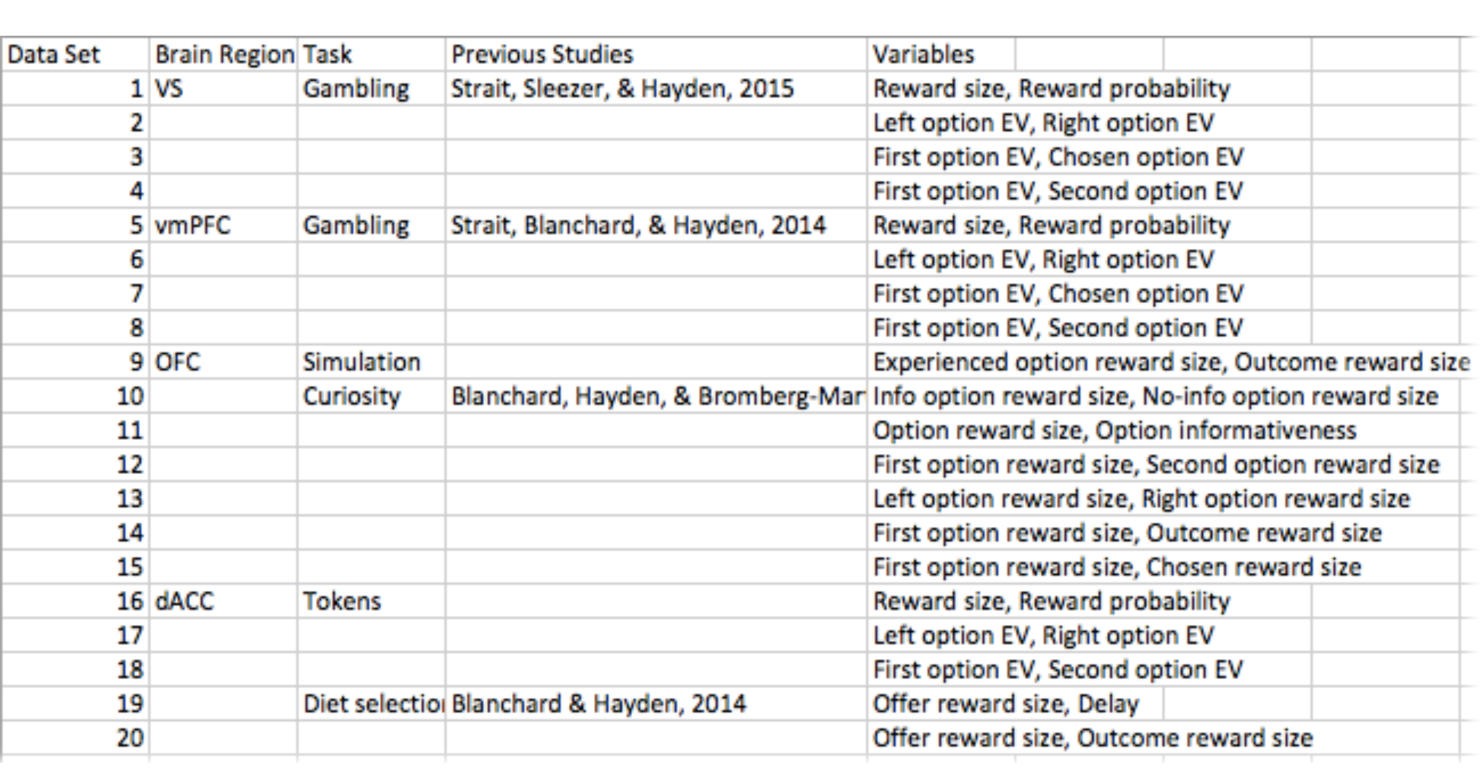
Analyses performed.

### Estimating the proportion of no-selectivity (i.e., noise) neurons

One benefit of our method is that it allows us to estimate the proportion of no-selectivity neurons (that is, neurons whose responses are only noise and not driven by task variables) in our population. We know of no previous estimate for this parameter in frontostriatal regions. Conventional analysis methods make it impossible to know if undriven cells are truly not task-related, or whether more trials would allow such neurons to pass a significance threshold. We report these proportions in **Figure 6C**.

In analyses of physiological data, correlation is often used to detect relationships between tuning to two different variables. For example, correlations between regression weights can provide information about integrated vs. disjoint (i.e. multiplexed) coding schemes. One limitation of this approach is that it underestimates the true correlation, raising the possibility of Type II errors. Specifically, a correlation using the entire population will produce an estimate biased by neurons that are not task responsive.

Our method allows us to reduce the decisional weight of neurons that have a low probability of being part of the mixed-tuning component. For this reason, we expect correlations based on our method to be stronger than those estimated using conventional approaches (and also to be more accurate). Indeed, this is what we found. For all data sets with significant correlations (positive or negative), our method estimates a stronger correlation than the standard method (**Figure 7**).

**Figure 7.**
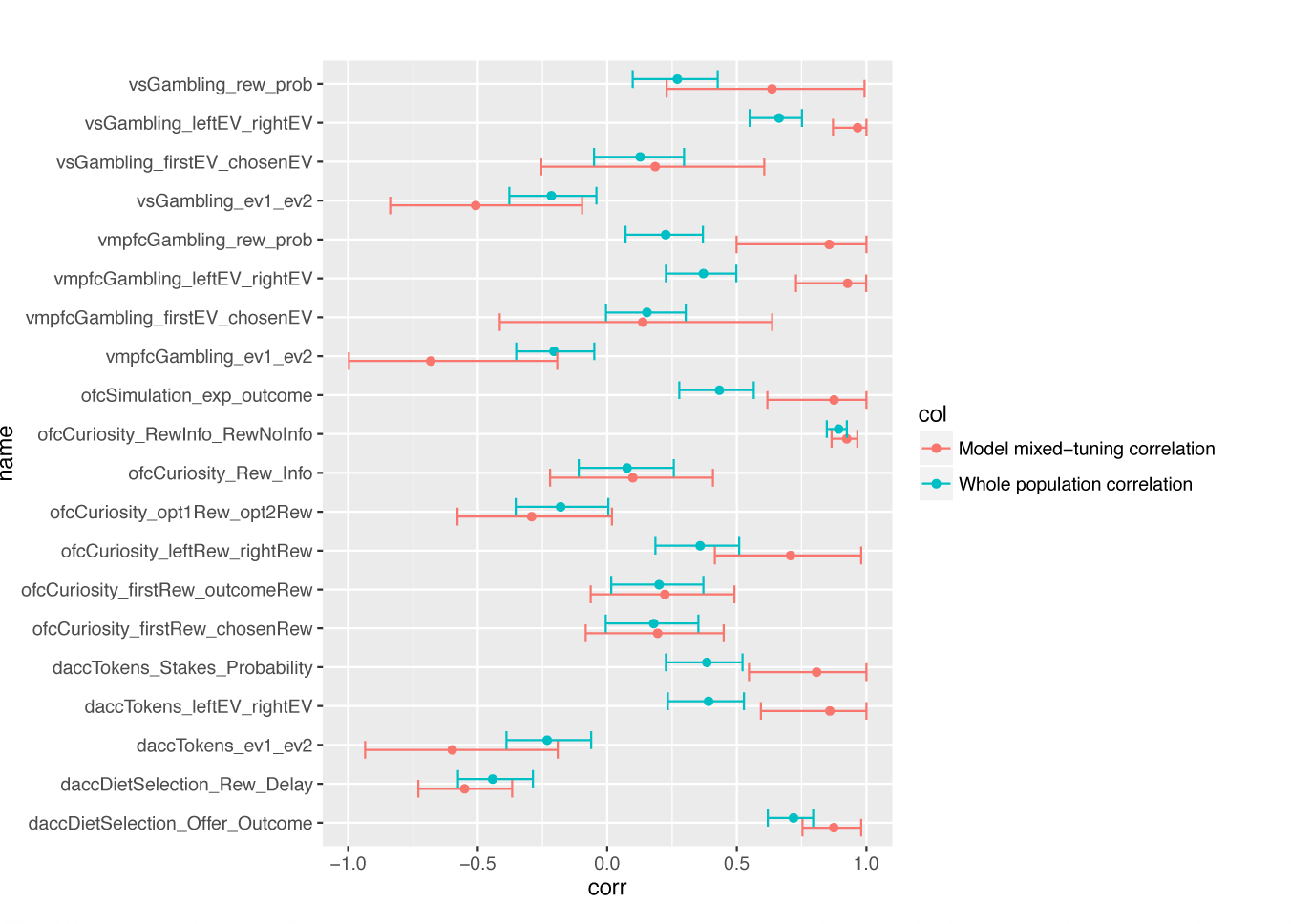
Illustration of measured correlations and confidence intervals using standard methods (green bars) and using our methods described here. In all cases with significant effects, measured correlations were more extreme (farther from zero) using our method.

### Public disposition of our code and data

All code relevant for using our analysis procedure is on github, along with instructions for use. All the data presented here are also available there. The address is https://github.com/TommyBlanchard/StanNeuronModelling

## DISCUSSION

Based on the tuning of each neuron in a sample, a population of cells can, in principle, be organized into functionally discrete clusters with categorically distinct response patterns. We developed a new statistical method that estimates the proportion of neurons that belong, respectively, to no selectivity (noise, firing unrelated to task parameters), pure selectivity (firing selective for a single task parameter), and/or mixed selectivity (firing selective for two task parameters simultaneously) categories. Reanalysis of our own data from previous studies reveals the presence of two discrete categories consistently for several brain areas and comparisons: one category containing task-insensitive cells (noise neurons) and one containing neurons sensitive to task variables. These findings raise the possibility that, for many variables of interest to neuroscientists, populations of neurons may be category-free (Raposo et al., 2014). Mixed selectivity has many convenient features, including scalability, flexibility, robustness to damage, and the lack of need for precise wiring (Rigotti et al., 2013; Fusi et al., 2016; Barak et al., 2013; Raposo et al., 2014). Our results are also consistent with a view in which ensemble responses provide the best description of neuronal decision-making and control systems, one that is largely independent of the properties of elemental cells (Mante et al., 2013; Stokes et al., 2013; Churchland et al., 2012; Kristan and Shaw, 1997; Ma et al., 2014; Hyman et al., 2012).

One unexpected finding of our analysis is that many neurons fall into the category of no-selectivity neurons – that is, the algorithm believes it has enough data to say with confidence that they are non selective for task variables, and not simply lacking enough data. Of course, these neurons presumably have some function. For example, our tasks were all oculomotor; perhaps they were purely manual modality cells. In any case, identifying the functional role of these cells is an important goal for future studies. One immediate benefit of this finding is that, by assessing the amount of noise associated with a given neuron, we can more accurately estimate population parameters, such as the correlations between tuning parameters in neuronal data. In any case, the identification of such cells presents a great opportunity to improve estimates of parameter correlations among driven cells in a population.

Classification of neurons by their functional properties is an appealing way to describe large and heterogeneous datasets. However, standard approaches, such as k-means clustering, require additional ad-hoc analyses in order to determine whether a given clustering is more likely than chance. Our Bayesian method tackles this problem directly and using minimal assumptions. One recent formal method, the PAIRS test provides an important tool and important evidence for non-categorical tuning in one region (Raposo et al., 2014). Our model extends on this earlier work in four ways. First, in that work, the null hypothesis is no categories, and thus a result of “no categories” provides a somewhat limited conclusion, as failures to reject the null do. Our Bayesian approach gets around this limitation. Second, our method is flexibly able to deal with any relationship between the tuning properties of neurons to different variables. Thus it is more general and can deal with more variety of population structures. Third, our method identifies, with confidence levels, information about what neuron types compose a population. Finally, the PAIRS test does not deal in a principled way with the possibility that only a subset of neurons measured may be task-relevant; that is, it ignores a possible no-selectivity cluster. Nonetheless, the PAIRS test represents an important and valuable tool for the field, one that is complementary to ours. The results of that test in rodent parietal cortex presents an important challenge to the idea of categorical tuning, one that our results echo.

It is very common to perform statistical hypothesis testing and then do further analyses based on the results of such tests. Such a practice is ill-advised. By using what is ultimately an arbitrary a statistical significance threshold (normally p<0.05) to categorize data, one is imposing a hard categorical boundary in the face of what is in reality graded statistical support for the categorization. Further analyses on such data necessarily produces Type II errors, and can, in some cases produce type I errors (Maxwell and Delaney, 1993; MacCallum et al., 2002). The standard approach likely also introduces a potential non-independence error: by removing cells that do not achieve a given p-value, the remaining cells are certain to be non-representative of the population in an anti-conservative direction: examination of only cells that pass the threshold can give results that appear, incorrectly, to be statistically strong. A similar problem is known in neuroimaging as “double-dipping,” which selects the brain region of interest with one statistical test and subsequently conducts non-independent analysis on the selected region (Kriegeskorte et al., 2009). Our methods offer a new way around this problem. Finally, in cases where the no-selectivity neurons are scientifically important, a conservative cutoff (like p<0.05) is also likely to overestimate the proportion of no-selectivity cells in the population, thus leading to false positives (Maxwell et al., 1984; Humphreys and Fleishman, 1974).

It is important to describe several limitations of our modeling approach in order to define the boundaries of where our method is most usefully applied and to point towards further extensions of the general approach. First, the model assumes a specific form of mixture components (non-selective, pure-selective, and mixed-selective). If there were a population that did not fit into these categories - for instance, two distributions with different orientations - then the population would not be well modeled by our existing framework. Similarly, if the distributions did not have zero mean or were not unimodal, the present method would not be appropriate. (Our data were z-scored). This limitation may not be as bad as it seems. Any such data could be incorporated in principle into a mixture model like ours by mathematically defining and including the corresponding distribution with its associated free parameters. For instance, numerous distributions with free orientations and means could be used in situations where there are few specific hypotheses about how neural responses are likely to behave. In this way, our approach is an instantiation of a more general technique that is increasingly common, Bayesian data analysis, where a family of hypothesized generative models may be written down, and the data can be used to optimally infer how likely each model or parameter value is. This family of extensions of our model which focuses on modeling the underlying components of the data is likely to yield deep insights into the organization of neural systems.

## MATERIALS AND METHODS

### Model

The model was implemented in the programming language Stan (Carpenter et al., 2016). Code is available on Github and distributed under a GPL3 license Address:https://github.com/TommyBlanchard/StanNeuronModelling

We ran MCMC for 5000 steps and 5 chains. Convergence was assessed using visual analysis of trace plots and confirmed by the Gelman and Rubin convergence diagnostic (called 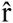, Gelman and Rubin, 1992). The variable 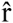 (which specifically measures the ratio of variance across different simulations) was below 1.05 for all variables for all data shown, indicating that the numerical methods used in the analysis accurately recovered the true posterior parameter values.

To conduct this analysis, we must assume priors on the likely values of all parameters. The priors were chosen to have as little influence as possible on the results, meaning that we let the data “speak for itself.” We used non-informative (Jeffrey’s) priors on the mixture weights and a LKJ prior on the correlation matrices that is uniform over all such matrices (Gelman & Hill, 2006). The scale parameters used half-Cauchy priors that permit a large range of possible values with very little penalty.

### Generation of physiological data

All data come from previously published studies; detailed methods are given in the appropriate citation. The following is a brief summary.

#### Surgical procedures

All animal procedures were approved by the University Committee on Animal Resources at the University of Rochester and were designed and conducted in compliance with the Public Health Service’s Guide for the Care and Use of Animals. Male rhesus macaques *(Macaca mulatta)* served as subjects.

#### Recording sites

We approached vmPFC, VS, OFC, and dACC through standard recording grids (Crist Instruments). Images of the recording sites and tasks can be found in the appropriate papers. We defined vmPFC as Area 14, lying within the coronal planes situated between 29 and 44 mm rostral to the interaural plane, the horizontal planes situated between 0 and 9 mm from the brain’s ventral surface, and the sagittal planes between 0 and 8 mm from the medial wall. We defined VS as NAcc core lying within the coronal planes situated between 28.02 and 20.66 mm rostral to interaural plane, the horizontal planes situated between 0 to 8.01 mm from ventral surface of striatum, and the sagittal planes between 0 to 8.69 mm from medial wall. We defined OFC as Area 13, lying within the coronal planes situated between 29.50 and 35.50 mm rostral to interaural plane, the horizontal planes situated between 0 to 6.00 mm from the brain’s ventral surface, and the sagittal planes between 6.54 to 13.14 mm from medial wall. We defined dACC as Area 24 (for a discussion of our decision to use this name, see Heilbronner and Hayden, 2016), lying within the coronal planes situated between 29.50 and 34.50 mm rostral to interaural plane, the horizontal planes situated between 4.12 to 7.52 mm from the brain’s dorsal surface, and the sagittal planes between 0 and 5.24 mm from medial wall.

#### Electrophysiological techniques

Single electrodes (Frederick Haer & Co., impedance range 0.8 to 4M Ω) were lowered using a microdrive (NAN Instruments) until waveforms between 1 and 3 neuron(s) were isolated. Individual action potentials were isolated on a Plexon system (Plexon Inc., Dallas, TX). Neurons were selected for study solely on the basis of the quality of isolation; we never pre-selected based on task-related response properties. All collected neurons for which we managed to obtain at least 300 trials were analyzed; no neurons that surpassed our isolation criteria were excluded from analysis.

#### Eye-tracking and reward delivery

Eye position was sampled at 1000 Hz by an infrared eye-monitoring camera system (SR Research). Stimuli were controlled by a computer running Matlab (Mathworks) with Psychtoolbox and Eyelink Toolbox. Visual stimuli were colored rectangles on a computer monitor placed 57 cm from the animal and centered on its eyes. A standard solenoid valve controlled the duration of juice delivery. The relationship between solenoid open time and juice volume was established and confirmed before, during, and after recording.

#### Behavioral tasks

Subjects performed in four different tasks with the same basic structure and one task with a somewhat different structure. For the neuronal recordings in vmPFC, subjects B and H performed the *risky choice task;* for VS, subjects B and C performed the *risky choice task;* for OFC study 1, subjects B and J performed the *curiosity gambling task;* for OFC study 2, subject B and H performed the *riskless choice task;* for dACC study 1, subjects B and J performed the *token risky choice task;* for dACC study 2, subjects B and J performed the *diet selection task.* All tasks made use of rectangles indicating reward amount and either probability or (in the diet selection task) delay. This method produces reliable communication of abstract concepts such as reward, probability, and delay to monkeys (see corresponding references, and also Hayden et al., 2010; Pearson et al., 2010; Blanchard et al., 2013; Blanchard and Hayden, 2015, for quantitative analyses demonstrating the robustness of these behavioral methods).

#### Risky choice task

Two offers, indicated by rectangles, were presented on each trial. Options offered a risky bet for liquid reward (there were safe offers as well, which were excluded from these analyses). Gamble offers were defined by two parameters, *reward size* and *probability.* The size of the blue or green portion of the rectangle signified the probability of winning a medium (mean 165 μL) or large reward (mean 240 μL), respectively. These probabilities were drawn from a uniform distribution between 0 and 100 %. The rest of the bar was colored red; the size of the red portion indicated the probability of no reward. On each trial, one offer appeared on the left side of the screen and the other appeared on the right. The sides of the first and second offer (left and right) were randomized by trial. Each offer appeared for 400 ms and was followed by a 600 ms blank period. After the offers were presented separately, a central fixation spot appeared and the monkey fixated on it for 100 ms. Following this, both offers appeared simultaneously and the animal indicated its choice by shifting gaze to its preferred offer and maintaining fixation on it for 200 ms.

#### Curiosity gambling task

A similarly structured gambling task, where gambles always carried a reward probability of 50 % and the size of a white bar in the center of each offer indicated *reward size* at 21 levels: from 75 to 375 μL in increments of 15 μL. Each trial, the monkey chose between an *informative* gamble (cyan; if chosen a cue 2.25 s before the potential reward would indicate if the monkey was about to win) and an *uninformative* gamble (magenta; if chosen the cue was replaced with an uninformative decoy; Fig. 2B). The stakes of both options and the order and side on which the informative option appeared were all randomized on all trials. Critically, the information was not revealed during the presentation of the cues, but only after the choice was made. Thus neural responses to the offers were not themselves reflective of the information.

#### Riskless choice task

This task was structured similarly to the other tasks, with the following exceptions: all cues were 100% valid. The first offer was one of five values [50 100 150 200 250] μL; the second offer was one of three values [150 175 200] μL. Half of trials (randomly selected by trial) were experienced (offer was given and choice of the offer would repeat it) and half were described (offer was indicated with a valid visual cue).

#### Token risky choice task

Another similarly structured gambling task, where gambles each had 2 potential outcomes, wins or losses in terms of “tokens” displayed onscreen as cyan circles. A small reward (100 μL) was administered concurrently with gamble feedback on each trial, regardless of gamble outcome. Trials in which the monkey accumulated 6 or more tokens triggered an extra “jackpot” epoch in which a very large reward (300 μL) was administered (**Fig. 2C**).

#### Diet selection task

Based on the famous foraging task of the same name. On each trial, a rectangle glided from top to bottom of screen at a rate such that it would take one second to traverse the screen. Fixation of the rectangle caused it to pause and begin shrinking at a constant rate. Successful maintenance of fixation led to a juice reward corresponding to the rectangle’s color [75 135 212 293 293/0] μL. (The symbol 293/0 indicates a gamble with 50% probability of 293 and 50% of zero). Widths indicated delay and were drawn from [2468 10] seconds.

## ACKNOWLEDGEMENTS

We thank Habiba Azab, Meghan Castagno, Giuliana Loconte, Marc Mancarella, Brianna Sleezer, Caleb Strait, and Maya Wang for help with data collection and organization. This work was supported by an R01 (DA037229) to BYH.

